# Virion-associated spermidine transmits with Rift Valley fever virus particles to maintain infectivity

**DOI:** 10.1101/2020.01.23.915900

**Authors:** Vincent Mastrodomenico, Jeremy J. Esin, Shefah Qazi, Oreoluwa S. Omoba, Brittany L. Fung, Maxim A. Khomutov, Alexander V. Ivanov, Suchetana Mukhopadhyay, Bryan C. Mounce

**Author notes:** To whom correspondence should be addressed: Department of Microbiology and Immunology, Loyola University Chicago, Stritch School of Medicine, 2160 S. First Ave. Maywood, IL 60153.

## Abstract

Viruses require host cell metabolites to productively infect, and the mechanisms by which viruses usurp these molecules is diverse. One group of cellular metabolites important in virus infection is the polyamines, small positively-charged molecules involved in cell cycle, translation, and nucleic acid synthesis, among other cellular functions. Polyamines also support replication of diverse viruses, and they are important for processes such as transcription, translation, and viral protein enzymatic activity. Rift Valley fever virus (RVFV) is a negative-sense RNA virus that requires polyamines to produce infectious particles. In polyamine depleted conditions, noninfectious particles are produced that interfere with virus replication and stimulate immune signaling. Here, we find that RVFV relies on virion-associated polyamines to maintain infectivity. We show that RVFV replication is facilitated by any of the three biogenic polyamines; however, we specifically find spermidine associated with purified virions. Using a panel of polyamine homologs, we observe that virions can also associate with (*R*)-3-methylspermidine and norspermidine, though not with other less homologous molecules. Using polyamine reporter cells, we demonstrate that virion-associated polyamines transmit from one infected cell to another. Finally, we find that virions devoid of polyamines are unstable and cannot be supplemented with exogenous polyamines to regain stability or infectivity. These data highlight a unique role for polyamines, and spermidine in particular, in maintaining virus infectivity, a function not previously appreciated. Further, these studies are the first to identify polyamines associated with RVFV virions. Targeting polyamines represents a promising antiviral strategy, and this work highlights a new mechanism by which we can inhibit virus replication through FDA-approved polyamine depleting pharmaceuticals.

## Introduction

Rift Valley fever virus (RVFV) is a significant human and ruminant pathogen, associated with hemorrhagic fever and spontaneous abortion. While the virus is currently geographically restricted to Africa and the Middle East, the potential for spread globally is significant. Further, RVFV is a mosquito-borne virus, and chikungunya^1^ and Zika^2^ virus demonstrate that these viruses can spread globally and explosively. Both *Culex* and *Aedes* species of mosquitoes transmit RVFV^3–5^, though the breadth of vectors susceptible to RVFV is not fully understood. Fortunately, several vaccine candidates^6–8^ show promise in reducing transmission, including in animals. However, adverse events associated with these vaccines have limited their use, and the virus continues to present itself in frequent outbreaks^9–11^, infecting hundreds and severely impacting local economies. Thus, the development of improved vaccines or the identification of novel antiviral targets is essential to the treatment and prevention of RVFV.

As obligate intracellular pathogens, viruses rely on their host cells for the building blocks of replication. These building blocks include a variety of metabolites produced by the host cell. One set of these metabolites crucial to virus replication is the family of polyamines. Eukaryotic cells synthesize polyamines to support transcription, translation, and cell cycling^12–14^. The biogenic polyamines include putrescine, spermidine, and spermine, which are maintained at millimolar level within cells^15^ and readily interconvert within cells^16^. These molecules are carbon chains of increasing length with primary and secondary amine groups. At physiological pH, polyamines are positively charged, which facilitates nucleic acid interactions. In fact, upwards of 85% of polyamines are bound to nucleic acids (primarily RNA), proteins, or lipids to support cellular functions^17^. However, polyamines are dispensable for cellular homeostasis in non-transformed cells. Depletion of polyamines via the specific inhibitor difluoromethylornithine (DFMO) reduces cellular proliferation but is otherwise nontoxic^18^. In humans, chronic DFMO treatment has mild side effects and is used in the treatment of trypanosomiasis^19,20^. Additionally, diethylnorspermidine (DENSpm) is a nontoxic molecule that enhances polyamine catabolism by acetylating polyamines and promoting their export or degradation. Thus, while polyamines are crucial to cellular replication, organismal polyamine depletion is tolerable.

Early work demonstrated that a subset of viruses incorporate polyamines in virions, especially large DNA viruses like herpesviruses and vaccinia virus, which use polyamines to package their large dsDNA genomes^21–23^. In contrast, RNA viruses, with relatively smaller single-stranded genomes were poorly studied in the context of polyamine metabolism. Polyamines facilitate RNA virus replication, and the polyamine inhibitor DFMO reduces the replication of diverse RNA viruses, including alphaviruses, flaviviruses, enteroviruses, and bunyaviruses, both *in vitro* and *in vivo*^24,25^. Chikungunya and Zika viruses (CHIKV and ZIKV) rely on polyamines for genome replication and viral polyprotein translation^24^. Additionally, we have shown that polyamines enhance viral protease activity^26^, promote infectious particle production^27^, and support virus-cell binding^28^ in enteroviruses and bunyaviruses. The breadth of mechanisms by which polyamines support virus infection remain unknown but recent evidence suggests that different viruses utilize polyamines via different mechanisms.

The distinct structures of polyamines have different roles in cells and in viruses. For instance, spermidine copurifies with *E. coli* tRNAs^29^ and each biogenic polyamine has a distinct affinity for tRNA^30^. Spermidine is also used specifically in the genesis of the modified amino acid hypusine, which is important for translation^31,32^ and also crucial in the replication of some viruses^33,34^. Herpesviruses preferentially package spermidine and spermine but not putrescine in their virions^21^, though it is unknown why these polyamines are preferred. In contrast, chikungunya virus polymerase is stimulated by polyamines and was equally stimulated by putrescine, spermidine, or spermine^24^, suggesting that some viral processes may be insensitive to polyamine identity. Similarly, phage T7 polymerase is stimulated by spermidine and a variety of polyamines not synthesized in eukaryotic or prokaryotic cells^13^.

Polyamines are crucial to RVFV infection, and we demonstrated that polyamine depletion reduces RVFV titers and leads to the production of noninfectious particles that interfere with virus replication^27^. Precisely how polyamines function in RVFV infection remains unclear, however. Here, we investigated whether RVFV relied on specific polyamines for replication. We observe that RVFV replication is supported by any of the biogenic polyamines as well as cadaverine and norspermidine, two bacterially-synthesized polyamines. We find that these polyamines support infectious particle production. We considered that polyamines may be associated with RVFV virions and measured them via fluorometric assay and thin layer chromatography. We identify spermidine within purified virions and that single-carbon modifications of spermidine can also associate with virions. We finally show that virion-associated spermidine enhances viral particle infectivity and that exogenous polyamines applied to virions cannot restore their infectivity. In sum, polyamines, spermidine in particular, are crucial to RVFV due to their association with the virion which maintains infectivity.

## Results

### Rift Valley fever virus is sensitive to low concentrations of biogenic polyamines

Bunyaviruses are sensitive to polyamine depletion mediated either by DFMO or DENSpm, and replenishing polyamines exogenously fully rescues replication. To determine if specific polyamines enhance virus replication, we depleted Huh7 cells of polyamines using 1 mM DFMO, infected at multiplicity of infection (MOI) of 0.1 plaque-forming units (pfu) per cell with RVFV strain MP-12 and then titrated the biogenic polyamines putrescine, spermidine, and spermine at the time of infection. After 48h, virus was collected and titered by plaque assay on Vero-E6 cells. We observed that DFMO significantly reduced viral titers compared to untreated samples, not supplemented with DFMO or exogenous polyamines (Figure 1, dashed line “DFMO” versus not treated, or “NT”). We further observed that viral titers remained suppressed until polyamine concentration passed 1 μM, which held for each of putrescine, spermidine and spermine (Figure 1A). We observed EC_50_ values of 3.6, 4.6, and 9.9 μM for spermine, spermidine, and putrescine, respectively. In fact, each polyamine rescued viral titers to levels that were not significantly different from untreated samples at 10 μM. We performed a similar analysis with distantly-related bunyavirus La Crosse virus (LACV) and observed similar results: all three biogenic polyamines supported infection in the micro-molar range (Figure 1B) and no polyamine was favored when supplemented to DFMO-treated cells.

**Figure 1.**
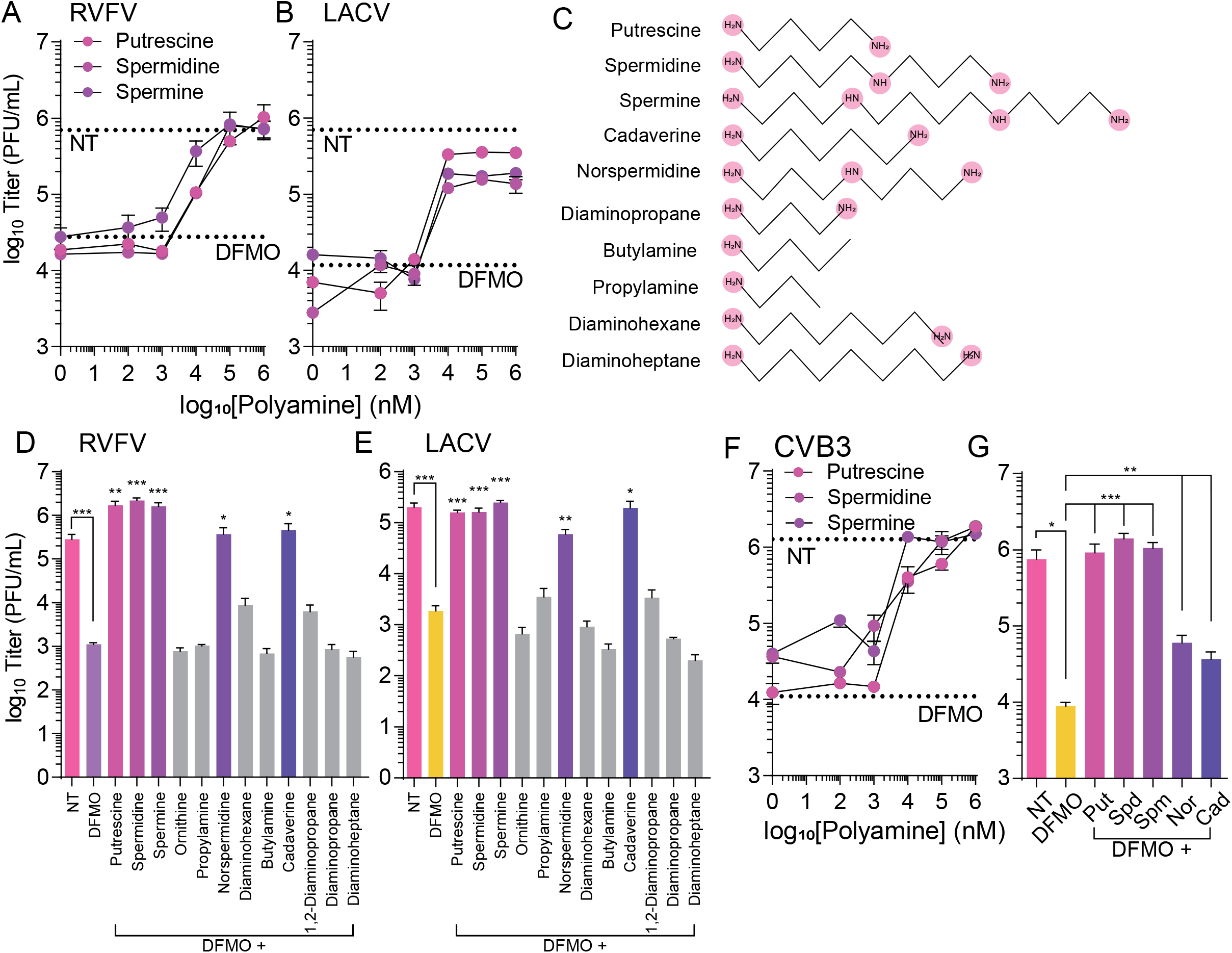
Biogenic and non-biogenic polyamines support RVFV replication. Huh7 cells were treated for four days with 1 mM DFMO and then infected with (A) RVFV or (B) LACV at MOI 0.1. Putrescine, spermidine, and spermine were added directly to the media at the time of infection at the concentration listed. Titers were determined by plaque assay at 48 hpi. (C) Chemical structures of polyamines analyzed for their ability to support viral infection of (D) RVFV and (E) LACV. Cells were treated as in (A) and (B) but at the time of infection, cells were supplemented with 10 μM polyamine as indicated. Titers were determined at 48 hpi. (D) Huh7 cells were treated as in (A) and subsequently infected with CVB3 at MOI 0.1 for 24h. Titers were determined by plaque assay. (G) CVB3 infections were treated with 10 μM putrescine (put), spermidine (spd), spermine (spm), norspermidine (nsp), and cadaverine (cad). Titers were determined at 24 hpi. Error bars represent one standard error of the mean. *p<0.05, **p<0.01, ***p<0.001 by two-tailed Student’s T-test. Comparisons in (D), (E), and (G) are DFMO versus treatment or as indicated.

### Polyamines not synthesized in eukaryotes also support viral replication

The biogenic polyamines are found in all eukaryotic cells examined. Given the relatively simple structure of polyamines, consisting of carbon chains with primary and secondary amines, we hypothesized that additional non-biogenic molecules with similar structures may also enhance virus replication. To test this, we treated Huh7 cells with 1 mM DFMO, infected with RVFV, and supplemented cells with an array of polyamines at 10 μM at the time of infection (a selection of structures shown in Figure 1C. All polyamines except putrescine, spermidine, and spermine are not synthesized in eukaryotic cells and are non-biogenic). When we measured viral titers at 48 hpi, we observed that biogenic polyamines fully rescued viral titers; however, we also observed that the polyamines cadaverine (1,4-diaminopentane, one carbon longer than putrescine) and norspermidine (one fewer carbon than spermidine), two polyamines not synthesized by eukaryotic cells, rescued viral titers to nearly equivalent levels as the biogenic polyamines (Figure 1D). Interestingly, we observed that no other non-biogenic polyamines enhanced titers beyond DFMO treatment levels, despite their structural similarity. Of particular interest, butylamine, which is similar to putrescine but lacks an amino group, failed to enhance replication. Additionally, elongating the putrescine carbon chain to six (diaminohexane) or seven (diaminoheptane) carbons or shortening the chain to three carbons also eliminated enhancement of virus replication. Again, we tested this set of polyamines with LACV and observed that cadaverine and norspermidine again enhanced viral titers, while all other compounds did not, suggesting conservation in polyamine usage between these two bunyaviruses (Figure 1E).

Finally, we used an enterovirus model of infection, Coxsackievirus B3 (CVB3) to test its sensitivity to distinct polyamines. As with RVFV and LACV, we treated cells with DFMO and supplemented with polyamines at the time of infection. When we titrated the three biogenic polyamines, we observed enhancement of replication at 1 μM of either putrescine, spermidine, or spermine (Figure 1F), similar to RVFV and LACV. We also tested whether CVB3 could utilize cadaverine or norspermidine for replication. When we supplemented DFMO-treated cells with either of these compounds, however, neither cadaverine nor norspermidine enhanced replication (Figure 1G). These data suggest that distinct virus families may rely on distinct polyamine structures for optimal replication.

### Polyamines maintain RVFV specific infectivity

We previously demonstrated that polyamine depletion limits virus replication via the generation of noninfectious particles. To test whether the biogenic polyamines also maintain specific infectivity of RVFV, we measured the ratio of genomes to infectious virus, as measured by plaque assay. We treated Huh7 cells with 1 mM DFMO prior to infection at MOI 0.1. At the time of infection, we added 100 μM putrescine, spermidine, or spermine. After 48 h, supernatant was collected for titration and RNA purification. RNA was reverse transcribed and analyzed for RVFV genomes using virus-specific primers. We then calculated the genome-to-PFU ratio as a measure of specific infectivity. Similar to our previous work, polyamine depletion increased the ratio of genomes to PFU (Figure 2A), suggesting reduced specific infectivity. However, addition of putrescine or spermine returned this ratio to untreated levels and modestly increased specific infectivity (fewer genomes per PFU) with spermidine treatment. We performed a similar analysis for LACV and observed the same phenotype: DFMO treatment increases the genome-to-PFU ratios, while the biogenic polyamines reduce to untreated levels (Figure 2B).

**Figure 2.**
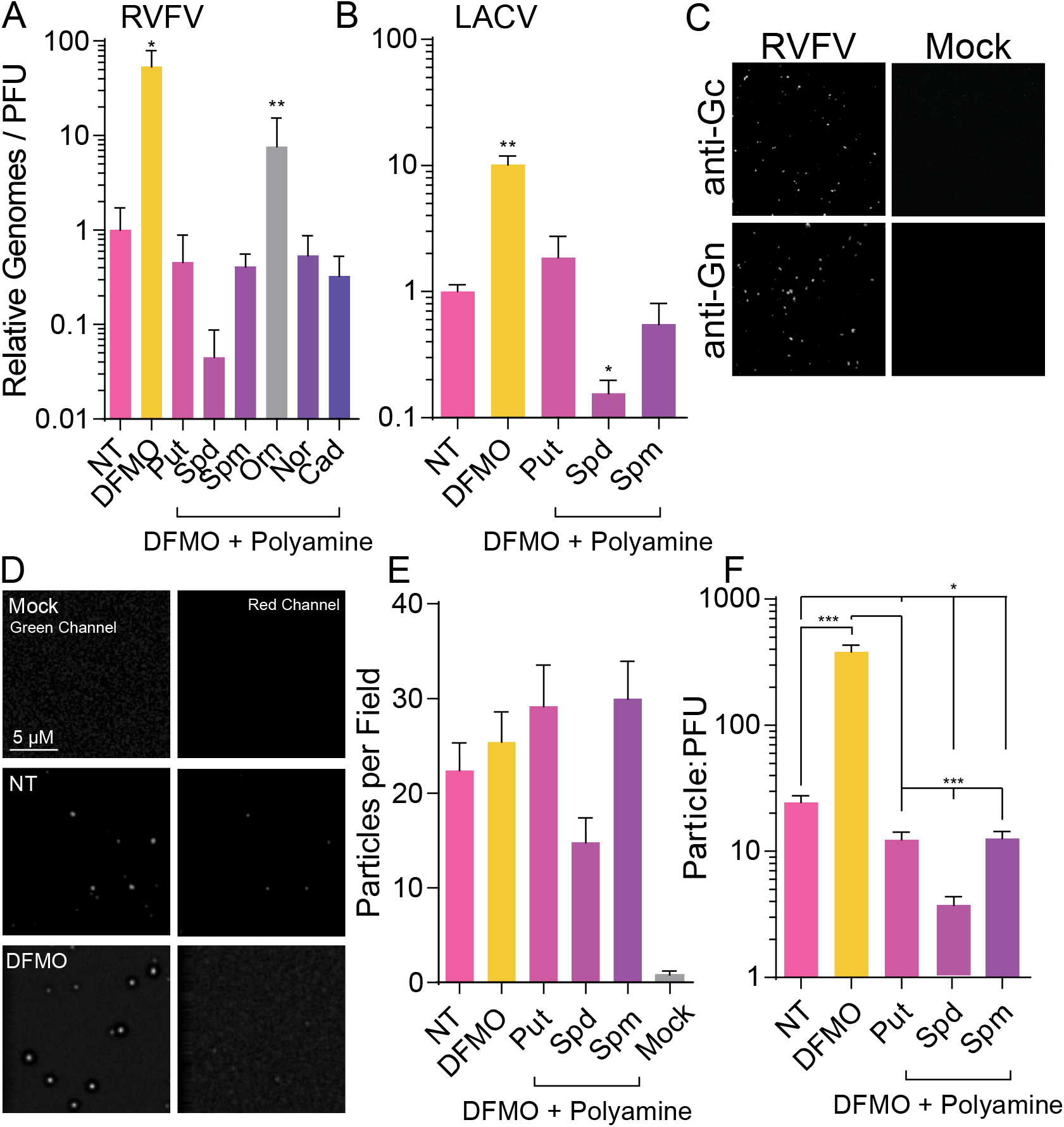
Biogenic polyamines enhance infectious particle production. Huh7 cells were treated for four days with 1 mM DFMO and subsequently infected at MOI 0.1 with (A) RVFV and (B) LACV for 48 h. Polyamines at 10 μM were added as indicated at the time of infection, and titers were determined by plaque assay and genome content determined by qPCR on reverse-transcribed viral RNA purified from cellular supernatant. Genome/PFU ratio was calculated by dividing the relative number of genomes by the viral titer. (C) Virus prepared as in (A) was spinoculated onto coverslips, fixed, and stained with anti-Gn or anti-Gc antibody. Mock-infected cell supernatant was similarly spinoculated as a control. (D) Virus prepared as in (A) was spinoculated and stained with anti-Gn and FITC secondary. Representative images from mock-infected, untreated infected, and DFMO-treated infected samples are shown in the green and red channels. (E) Particles from (D) were quantified with ImageJ and compared to viral titers to obtain (F) particle-to-PFU ratio. Images are representative from at least three independent preparations. Error bars represent one standard error of the mean. *p<0.05, **p<0.01, ***p<0.001 comparing untreated to treated conditions, unless otherwise specified, using a twotailed Student’s T-test.

Measuring genome-to-PFU is a surrogate for measuring the number of viral particles compared to the number of these particles that are infectious. To more accurately measure particle-to-PFU ratio, we used a method similar to Wichgers Schreur and colleagues to stain viral particles fluorescently^35^. We spinoculated virus on coverslips and stained for viral envelope glycoprotein Gc or Gn using specific antibodies and FITC-tagged fluorescent secondary antibody. To establish the assay, we tested both antibodies to ensure specificity and that we weren’t observing aberrations due to impurities on the coverslips or nonspecific staining. Using spinoculated virus derived from infected and uninfected cells, we observed dots corresponding to virus in only samples that were infected, no detectable signal was observed on slides spinoculated with samples from uninfected cells (Figure 2C). We next applied this method to virus derived from DFMO-treated cells as well as cells supplemented with various polyamines. Again, virus was spinoculated from mock-or virus-infected cell supernatant and stained with anti-Gn and FITC-tagged secondary. As a control, we also stained with a secondary antibody fluorescent in the red channel (TxRed). When we visualized the samples, we observed distinct puncta only in infected samples and not in mock samples (Figure 2D). We also observed no staining in the red channel, suggesting that we were again not observing impurities on the coverslips (Figure 2D, “Red channel”). We counted the number of dots using ImageJ and back-calculated the number of particles per mL of infected cell supernatant (Figure 2E). We used this number to calculate the particle-to-PFU ratio (Figure 2F). Supporting our genome-to-PFU ratios, we observed that untreated cells had a paticle-to-PFU ratio of approximately 20, and DFMO-mediated polyamine depletion increased this to >300. Thus, DFMO treatment increases the genome-to-PFU ratio, as well as the particle-to-PFU ratio, as measured by our two methods. To expand our rescue experiments, we similarly stained particles derived from infection of DFMO-treated and polyamine-supplemented cells. As with our genome-to-PFU ratio, we observe that addition of any of the biogenic polyamines returns the particle-to-PFU ratio to untreated levels, with a small, though significant, reduction in this ratio, suggesting that polyamines support RVFV infectivity.

### Polyamines are associated with the RVFV virion

We observe a change in specific infectivity (as measured by genome-to-PFU or particle-to-PFU), and we previously characterized that the physical and structural properties of virions produced with or without polyamines are indistinguishable. We next considered that polyamines might be associated with RVFV virions themselves. In fact, polyamines are found in the virions of several DNA viruses and a subset of RNA viruses. To this end, we used a fluorometric assay, which directly measures polyamine content in cells. We generated virus in Huh7 cells left untreated or treated with 1 mM DFMO by infecting at MOI 0.1 for 48 h. After 48 h, we purified virus by sucrose cushion ultracentrifugation, resuspended the viral pellet in PBS, and analyzed polyamine association. As a control, we used mock-infected supernatant and performed all steps in tandem with virus-infected cell supernatant. As expected, our purified mock-infected supernatant had no signal (Figure 3A). Similarly, CVB3-infected cell supernatant exhibited no detectable signal above background, as expected^23^. As a positive control^22^, we observed detectable levels of polyamines in purified vaccinia virus (VACV), and this signal returned to background levels when virus was derived from DFMO-treated conditions. When we tested RVFV, we observed signal above background, though not as intense as VACV, and this signal was depleted when virus was derived from DFMO-treated cells. These results suggest that purified RVFV virions are associated with polyamines.

**Figure 3.**
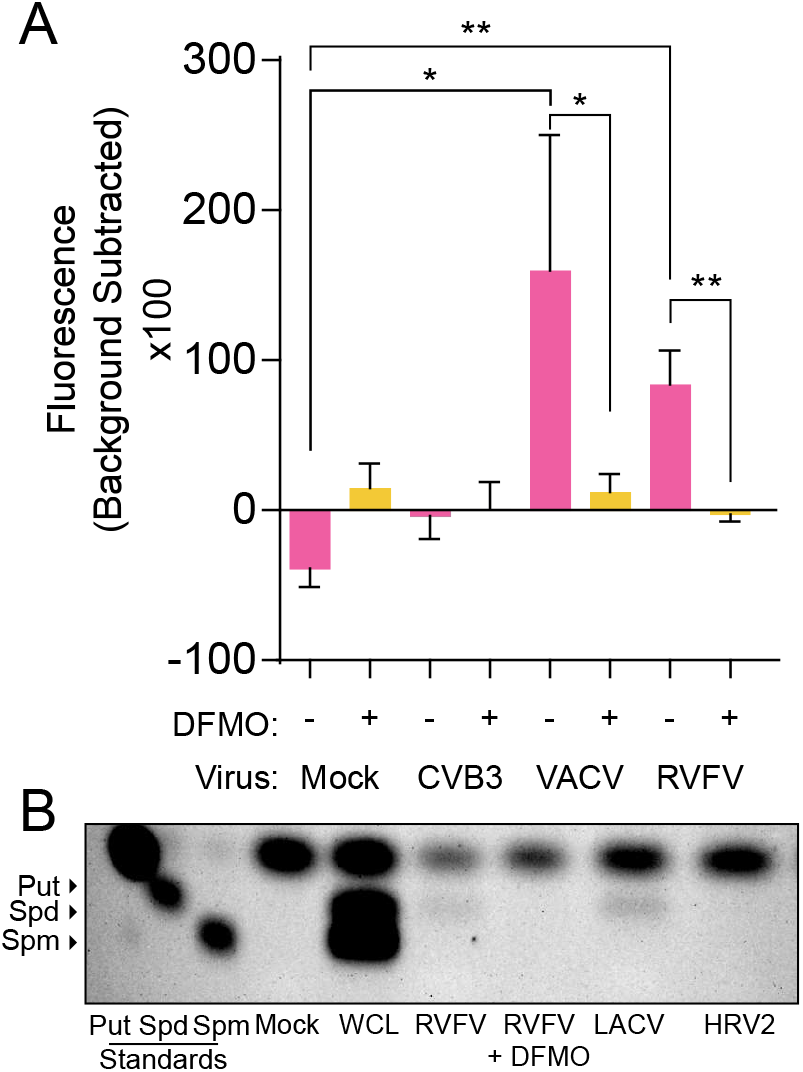
Polyamines associate with RVFV virions. Huh7 cells were treated for four days with 1 mM DFMO and infected at MOI 0.1 with CVB3, VACV, RVFV, LACV, or HRV2. At 48 hpi, cellular supernatant was collected and virus pelleted through a sucrose cushion prior to analysis by (A) fluorometric assay analyzing total polyamine content or (B) thin layer chromatography. Individual polyamines are indicated as put (putrescine), spd (spermidine) and spm (spermine). Error bars represent standard error of the mean. *p<0.05, **p<0.01 by two-tailed Student’s T-test comparing groups as indicated. Chromatogram displayed is representative of n=3 independent experiments.

Importantly, the fluorescent polyamine assay does not distinguish between polyamines in the cell; thus, this assay could not dictate which specific polyamine is associated with virions. In order to identify the polyamine(s), we purified and concentrated RVFV from Huh7 cells (approximately 10^6^ PFU total) as above, labeled polyamines via dansylation, and resolved the dansylated polyamines via thin layer chromatography (TLC). When we analyzed mock-infected cell supernatant, we observed no polyamines, as expected (Figure 3B). The whole cell lysate (WCL) from cells infected with RVFV contained robust amounts of spermidine and spermine, though little putrescine was detected. Interestingly, purified RVFV virions exhibited a distinct band corresponding to spermidine, and this band was lost when virus was purified from DFMO-treated cells. LACV also showed virion-associated spermidine, while human rhinovirus serotype 2 (HRV2, enterovirus distantly related to CVB3) had no detectable polyamines, as expected. These data again detect virion-associated polyamines, which we have identified as spermidine.

### Polyamines interconvert upon replenishment of DFMO-treated cells

We next considered whether we could deplete polyamines from cells, replenish with individual biogenic polyamines and detect these species in the RVFV virion. To this end, we treated Huh7 cells with 1 mM DFMO, infected with RVFV at MOI 0.1, and added 10 μM putrescine, spermidine, or spermine individually at the time of infection. As a control, we added ornithine, the polyamine precursor. As before, we purified and concentrated virions and analyzed polyamine content by thin layer chromatography. As expected, virions purified from DFMO-treated cells exhibited no polyamines; however, polyamine supplementation resulted in detectable virion-associated polyamines (Figure 4A). Interestingly, both putrescine and spermine supplementation led to a detectable level of virion-associated spermidine. Given that DFMO blocks the production but not the interconversion of polyamines, we considered that supplemented putrescine and spermine might generate spermidine through the actions of spermidine synthase (SMS) or polyamine oxidase (PAOX) with spermidine/spermine acetyltransferase (SAT1). We measured polyamine levels in DFMO-treated cells that were supplemented with the polyamines and observed that with putrescine, spermidine, or spermine supplementation, spermidine was abundant on our TLC (Figure 4B). Thus, the biogenic polyamines rapidly interconvert and specifically spermidine is virion associated in this polyamine milieu.

**Figure 4.**
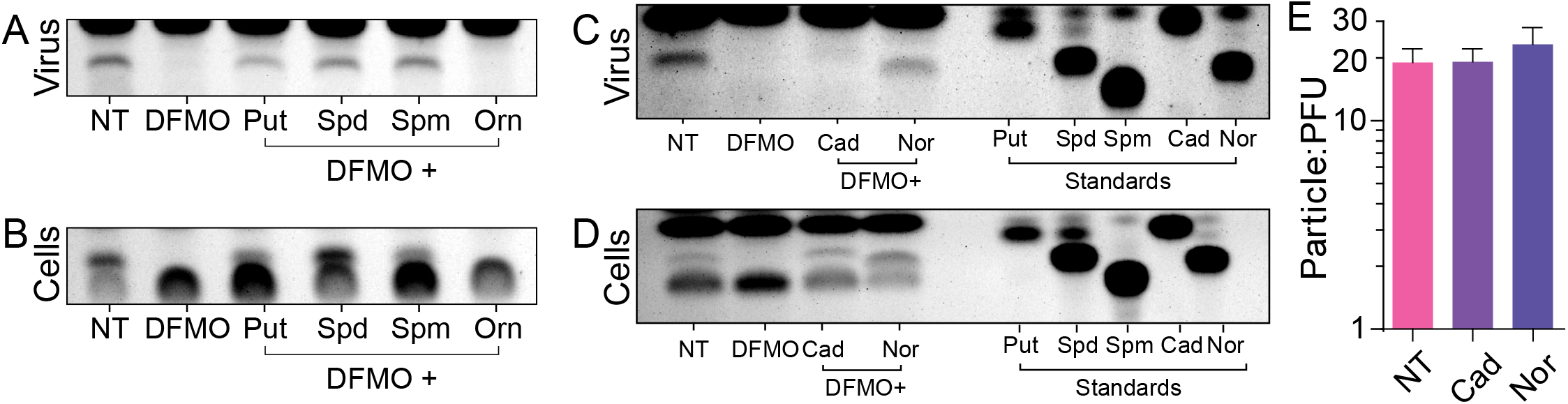
Polyamines interconvert upon replenishment of DFMO-treated cells. Huh7 cells were treated for four days with 1 mM DFMO and infected with RVFV at MOI 0.1 for 48 hpi. Polyamines were added to the cells at the time of infection. At 48 hpi, cells were collected and cellular supernatant virus purified for polyamine extraction. Polyamines were then visualized in (A) purified virions and (B) cells by thin layer chromatography. (C) Cells were treated and infected as in (A) but were supplemented with cadaverine (cad) or norspermidine (nor). Polyamine content of (C) purified virions and (D) cells was analyzed by thin layer chromatography. Chromatograms are representative of three independent experiments. (E) Virus prepared as in (C) were spinoculated onto coverslips, stained with anti-Gn antibody, and viral particles quantified. Particle counts were compared to titers to obtain the particle-to-PFU ratio. Error bars represent one standard error of the mean. No significant differences were determined by two-tailed Student’s T-test.

While eukaryotic cells can interconvert the biogenic polyamines, no description of their ability to interconvert cadaverine or norspermidine has been reported. Thus, we tested whether supplementation of these polyamines, which rescues viral titers, can support polyamine packaging. To this end, we generated and purified virus from DFMO-treated cells supplemented with 10 μM cadaverine or norspermidine and measure virion-associated polyamines by TLC. Curiously, we detected bands near the retention factor (Rf) of spermidine, though not precisely at spermidine’s Rf (Figure 4C). Our standards (Figure 4C, right) suggest that norspermidine is, in fact, associated with RVFV virions. However, the band in the cadaverine lane was faint and migrated slightly higher in the chromatogram. This band could be N(3-aminopropyl)cadaverine, which is a single carbon longer than spermidine, though this molecule has not been described to be synthesized in human cells. We checked whether we could detect this species in cells (Figure 4D), and we can in fact a molecule that runs as N(3-aminopropyl)cadaverine would. However, we also observe that norspermidine is not interconverted into other detectable polyamine species. In sum, however, it appears that polyamines within a limit can replace spermidine for RVFV virions.

### Methylated spermidine supports RVFV replication and is virion-associated

Polyamines rapidly interconvert, as observed (Figure 2B) and previously described. Methylated spermidine and spermine are not good substrates for acetylation by spermidine/spermine acetyltransferase and their interconversion is limited. (*R*)-3-methylspermidine is afunctionally active and metabolically stable analog of biogenic spermidine. However, (*R*)-3-methylspermidine is a poor substrate for spermine synthase and SAT1^36^. Thus, we considered whether methylated spermidine could enhance viral replication in the absence of biogenic spermidine and if this modified polyamine could be virion-associated. To test this, we treated cells with DFMO and replenished with (*R*)-3-methylspermidine (Figure 5A) at the time of infection with RVFV. When we measured titers at 48 hpi, we observed a full rescue in titers, to a level similar to spermidine. We titrated (*R*)-3-methylspermidine after DFMO treatment and observed that concentrations around 10 μM were sufficient to fully rescue viral titers (Figure 5B), slightly higher than for spermidine (Figure 1A). Thus, methylated spermidine functions to support RVFV infection.

**Figure 5.**
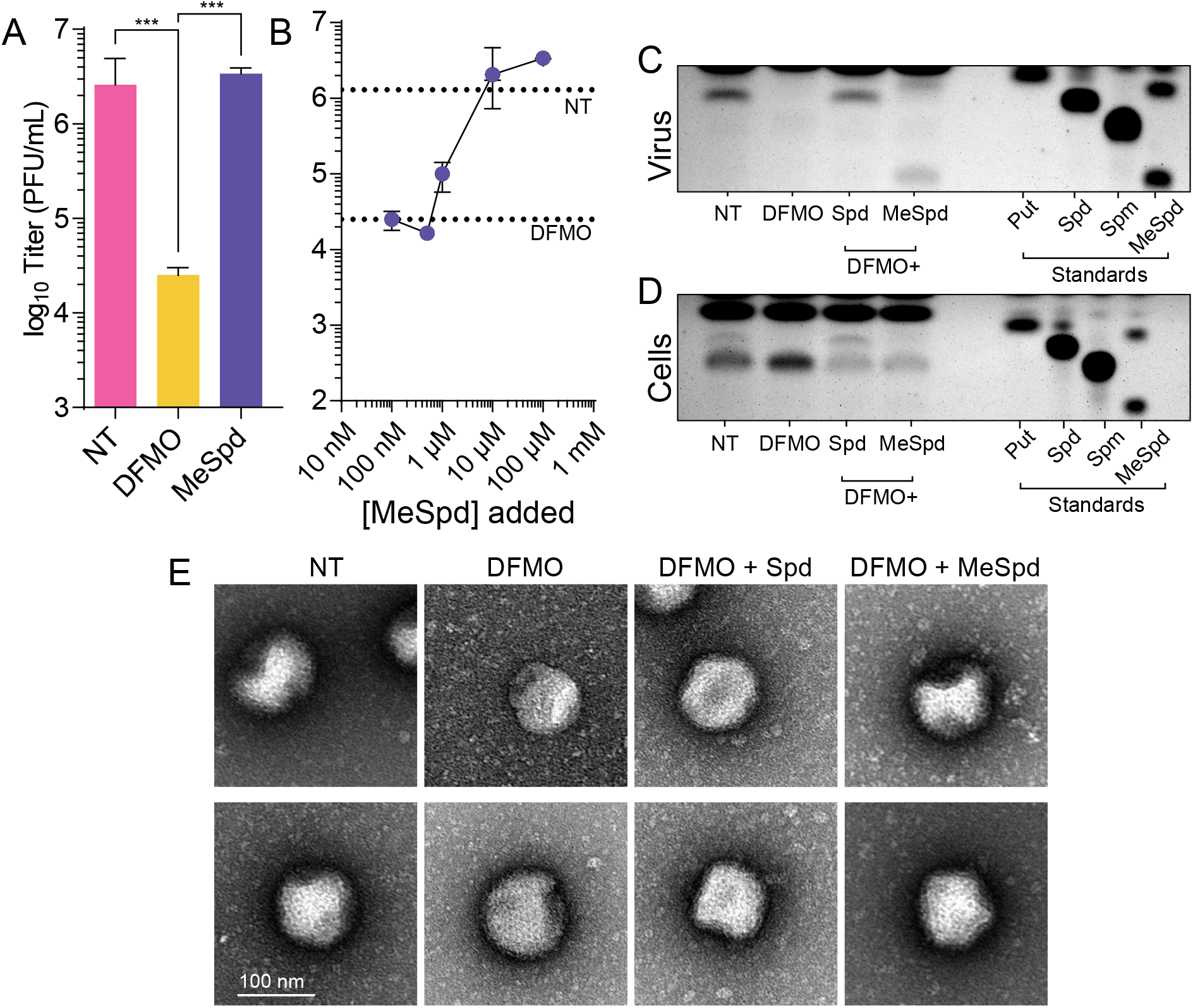
Methylated spermidine supports RVFV replication and is virion-associated. (A) Huh7 cells were treated with 1 mM DFMO for four days prior to infection with RVFV at MOI 0.1. Cells were supplemented with spermidine (Spd) and (*R*)-3-methylspermidine (MeSpd) at 10 μM. Viral titers were determined at 48 hpi. (B) Cells were treated and infected as in (A) but with increasing concentrations of (*R*)-3-methylspermidine. Titers were determined at 48 hpi. (C) Cells were treated and infected as in (A) and at 48 hpi, cellular supernatant was collected, virions purified, and polyamines extracted for analysis by thin layer chromatography. (D) Cells from (C) were collected and polyamine content analysed. (E) Representative electron micrographs of virus derived from untreated, DFMO-treated, or polyamine-supplemented conditions. Error bars represent one standard error of the mean.

To test if (*R*)-3-methylspermidine could associate with virions, we purified virions and visualized polyamines by TLC. As expected, we observed that RVFV was associated with spermidine; however, we could detect bands corresponding to (*R*)-3-methylspermidine as well (Figure 5D), suggesting that this polyamine is virion-associated. To confirm that 3-methylspermidine was not interconverted to the biogenic polyamines, we also performed TLC on the treated cells and observed no such interconversion (Figure 5E).

We previously showed that virions derived from DFMO-treated cells show no distinctions in their gross appearance by electron microscopy. To confirm this phenotype as well as to determine whether polyamine rescue with 3-methylspermidine could change virion morphology, we purified virions and examined them by electron microscopy. In untreated conditions, we observed numerous virions of expected size and with visible surface glycoproteins (Figure 5F). As previously described, DFMO treatment did not noticeably change virion appearance, and spermidine or 3-methylspermidine supplementation to DFMO-treated cells similarly had no discernible effect on virion appearance. Thus, polyamines do not appear to contribute to virion morphology.

### Polyamines are transmitted to naïve cells upon infection

Given that polyamines, specifically spermidine, are associated with RVFV virions, we considered that virus infection may transmit polyamines to newly-infected cells. We generated virus stock from untreated and DFMO-treated cells (Figure 6A) and then used these viruses to infect polyamine-sensitive luciferase reporter cells. These reporter cells consist of 293T cells transfected with an OAZ1 dual-luciferase construct. Briefly, OAZ1 transcript is sensitive to cellular polyamine levels: high polyamines result in enhanced stability and translation; low polyamines result in reduced stability and translation. With this construct, we can measure luciferase activity to indirectly measure polyamine levels in cells: polyamine levels directly correlate with firefly luciferase activity, which is normalized to renilla luciferase activity that is polyamine-independent. We treated these reporter cells with DFMO and observed a significant reduction in luciferase activity, corresponding to reduced polyamine levels. To these DFMO-treated reporter cells, we added 10 μM polyamines (equimolar mixture of the biogenic polyamines) and observed enhanced luciferase activity, demonstrating their responsiveness to polyamines. We also added purified RVFV virions (10^3^, 5×10^3^, and 10^4^ PFU), for which we observed a dose-dependent increase in signal (Figure 6B). When we added virus (10^4^ pfu) derived from DFMO-treated cells or when we purified supernatant from mock-infected cells, we did not observe an increase in luciferase activity, indicating that we are not observing a cellular response to infection or aberrantly purifying polyamines from cellular supernatant. In sum, these data suggest that RVFV can transmit polyamines upon infection.

**Figure 6.**
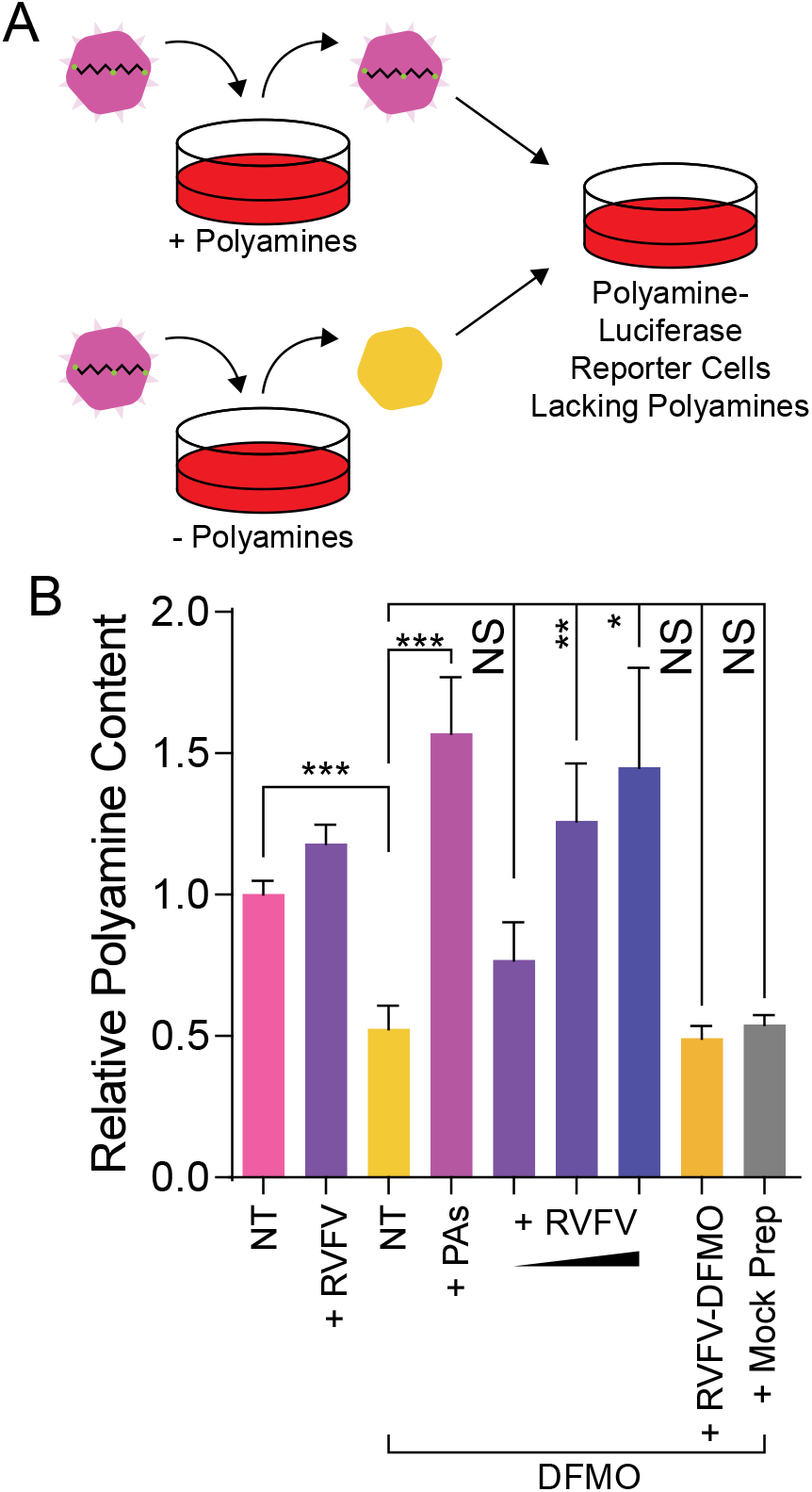
Polyamines are introduced to target cells upon infection. (A) Schematic of experimental setup. (B) 293T cells were treated with 1 mM DFMO for four days and subsequently transfected with a polyamine-sensitive dual-luciferase construct. Cells were subsequently left not treated (NT), supplemented with polyamines (a 100 μM mix of putrescine, spermidine, and spermine), or infected with purified RVFV as indicated. Luciferase activity was measured 24h later to calculate the relative polyamine content. RVFV-DFMO is RVFV derived from DFMO-treated, polyamine-depleted cells. Mock prep indicates treatment of transfected cells with supernatant purified as with virus purification. *p<0.05, **p<0.01, ***p<0.001, NS - not significant using a two-tailed Student’s T-test with comparisons as indicated.

### RVFV particles devoid of polyamines rapidly lose infectivity

Polyamines facilitate infection and polyamine depletion results in the genesis of non-infectious particles. Additionally, we can detect spermidine (and highly similar non-biogenic polyamines) in purified RVFV. We hypothesized that these non-infectious particles may be due to a decline in viral infectivity from a lack of virion-associated polyamines. To test this, we generated virus from untreated and DFMO-treated Huh7 cells and incubated the cell-free supernatant at 37°C for 24h, taking samples at regular intervals to titer. We observed that virus derived from untreated cells slowly declined in titer (Figure 7A), resulting in a modest reduction in titers over 24h. The calculated half-life of the virus was approximately 29.4h. In contrast, virus derived from DFMO-treated cells rapidly lost infectivity, with titers dropping by about 90% within 24h and a half-life of approximately 14.4h. Thus, polyamine-depleted cells generate virus that rapidly loses infectivity.

**Figure 7.**
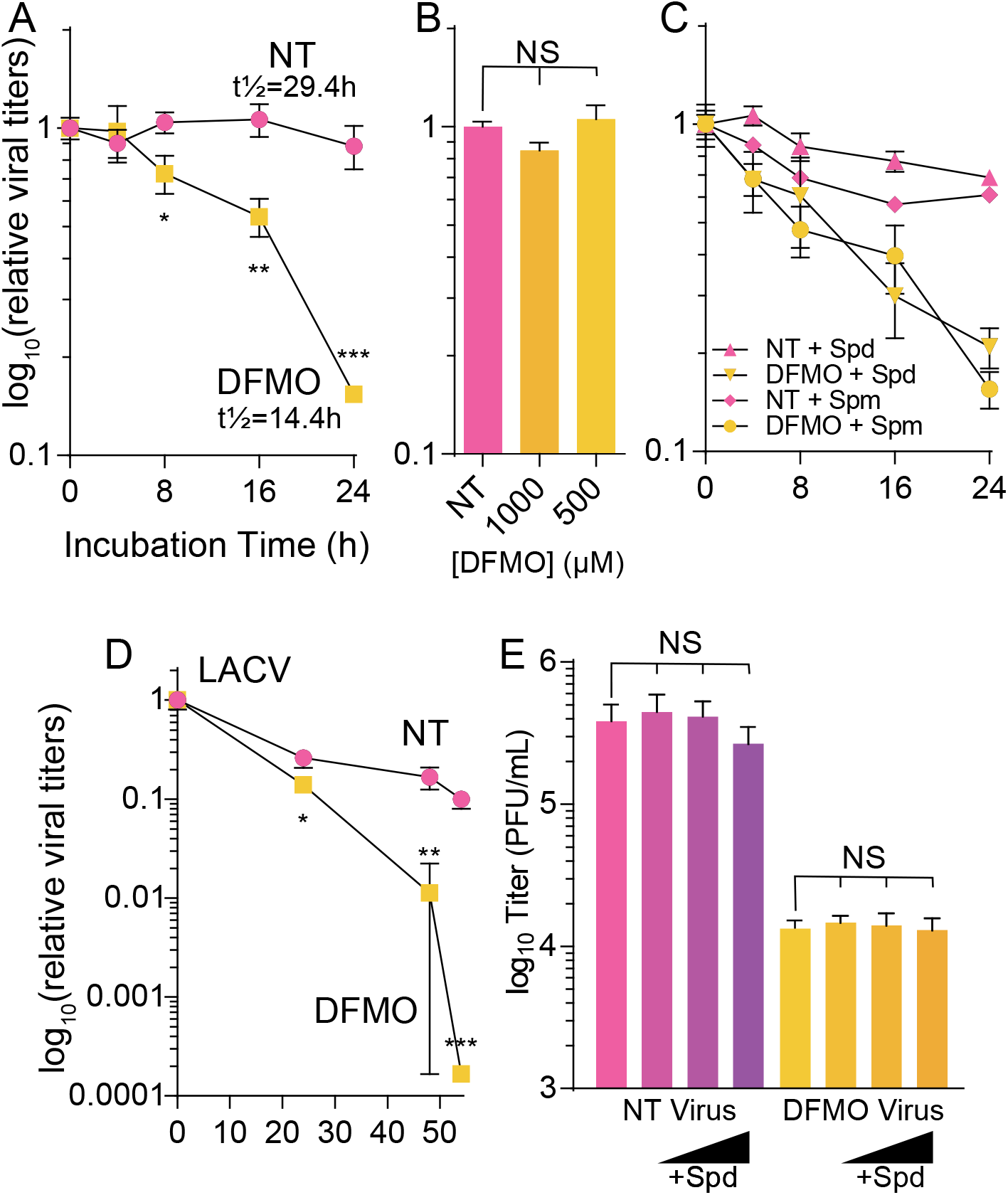
Polyamines maintain infectivity of bunyavirus particles. (A) RVFV derived from untreated or DFMO-treated cells was incubated at 37°C for the indicated time, when viral titers were determined by plaque assay. (B) RVFV stock virus was incubated with increasing doses of DFMO for 24h at 37°C. (C) Virus derived as in (A) was incubated with exogenous spermidine (Spd) or spermine (Spm) at 37°C for the indicated time prior to titering by plaque assay. (D) LACV derived and treated as in (A) was incubated at 37°C and titered by plaque assay. (E) RVFV was incubated with increasing doses of exogenous spermidine (Spd) for 24h at 37°C prior to plaque assay. *p<0.05, **p<0.01, ***p<0.001, NS not significant by two-tailed Student’s T-test comparing not treated (NT) conditions to DFMO.

We first considered that DFMO itself was destabilizing RVFV virions. Thus, we incubated virus with 1 mM or 500 μM DFMO for 24h at 37°C. We observed that DFMO itself did not destabilize the virus, as titers were equivalent between untreated and DFMO-treated viruses (Figure 7B). Because spermidine is found in association with purified RVFV virions, we next hypothesized that polyamines themselves may stabilize virions. To this end, we incubated virus generated from untreated or DFMO-treated cells with 10 μM spermidine or spermine and incubated at 37°C for 24h. We observed decay of the virus derived from DFMO-treated cells accelerated compared to untreated cells, with no difference regardless of polyamine treatment (Figure 7C). These data suggest that polyamines do not stabilize RVFV when added exogenously. Together, these data suggest that polyamines help to maintain infectivity, that this is not due to DFMO itself, and that exogenous polyamines cannot stabilize virions.

To extend these results to the related bunyavirus LACV, we similarly incubated virus derived from untreated and DFMO-treated conditions and measured infectivity over 24h. As with RVFV, we observed a steep decline in virus titers, though the effect was primarily at 48 hpi (Figure 7D). Thus, LACV exhibits similar sensitivity to losing infectivity when the virus is derived from DFMO-treated cells.

Finally, we considered whether we could potentially resurrect infectivity of our viral particles by incubating them with polyamines, specifically spermidine. To this end, we incubated virus from untreated and DFMO conditions with increasing doses of spermidine for 24h at 37°C. We observed no significant difference in titer from spermidine treatment (Figure 7E), suggesting that spermidine supplementation cannot rescue infectivity of the virions once they have lost infectivity.

## Discussion

As obligate intracellular pathogens, viruses rely on the host to provide metabolites for replication. The virion itself is composed of molecules derived from the host but directed for assembly and order by the viral genome. The genomic nucleotides, proteins’ amino acids, and envelope’s lipids originate from host metabolites. Here, we identify polyamines as an additional host-derived metabolite associated with bunyavirus virions. Previous reports have identified polyamines in the virions of herpesviruses^21^ and poxviruses^22^. However, other viruses do not appear to consistently incorporate polyamines into virions. For example, negligible amounts of polyamines are detected associated with poliovirus capsids, but rhinovirus 14 had a detectable amount, enough to neutralize approximately 16% of the genome^23^. Adenovirus-5, a DNA virus, also appears to incorporate small amounts of polyamines^37^. Thus, the packaging of polyamines and the role(s) of these packaged polyamines is not necessarily evolutionarily conserved. Additionally, the presence of these polyamines had not been examined in bunyaviruses.

We specifically observe that spermidine is associated with RVFV and LACV virions, despite cellular polyamine pools comprising primarily spermidine and spermine. Other viruses incorporate polyamines without specificity; for example, densoviruses have all three polyamines in purified virus^38^. Interestingly, herpesviruses package spermidine and spermine, with spermidine primarily associated with the envelope or tegument and spermine in the viral capsid^21^. Given that bunyaviruses like RVFV do not have a bona fide capsid, one could speculate that the viral envelope specifically associates with spermidine, given that both viruses have a lipid membrane component. The localization of polyamines in RVFV may further inform the function of these polyamines in the virion. Additionally, the mechanism by which virions incorporate spermidine but exclude the other polyamines is not known for this or other viruses. Whether spermidine incorporation is an active process by the virus or a product of RNA-, protein-, or lipid-spermidine interactions will be an important distinction to make.

Despite the specific association of spermidine with purified virions, we observe that any of the biogenic polyamines (putrescine, spermidine, and spermine) supports viral infection. Importantly, when treating cells with these polyamines, they interconvert and produce the full complement of cellular polyamines. In fact, we observe this in our system, and those cells replenished with any of the biogenic polyamines support virus infection and spermidine association with virions. Thus, the balance of polyamines is crucial to cellular homeostasis^16^ and virus replication. Interestingly, RVFV exhibits some flexibility in polyamine utilization, as we can detect molecules that are a single carbon longer or shorter than spermidine in virions. Interestingly, cells exhibit heterogeneity in their polyamine composition and, thus, if viruses differentially utilize polyamines, the cellular polyamine composition may alter infection and pathogenesis. Regardless, whether distinct polyamines function differently during RVFV infection remains to be fully understood.

We previously observed that polyamine depletion led to the accumulation of noninfectious viral particles that interfered with productive virus infection^27^. We were unable to find a physical distinction between infectious and noninfectious virions, however. These studies suggest that a component of the virus that may be maintaining infectivity is spermidine, as viruses lacking polyamines are more labile than viruses propagated in cells with polyamines. However, these results do not preclude that an additional polyamine-modulated factor may contribute to virion stability. In fact, polyamine depletion affects several cellular processes. Regardless, we observe that polyamine depletion generates viral particles lacking polyamines that rapidly lose infectivity. Future work will address the mechanisms behind this lability.

Diverse viruses rely on polyamines for their replication, and the diversity of viruses may rely on different polyamines for different processes. For example, Ebolavirus, a filovirus, utilizes polyamines for genome replication but hypusine, derived from spermidine, for protein translation^33,34^. As mentioned, herpesviruses package spermidine in viral envelope/tegument and spermine in capsids^21^. Whether these phenotypes are broadly shared is unclear, but understanding the mechanisms by which viruses utilize polyamines may highlight both evolutionarily conserved and divergent mechanisms. Importantly, however, the requirement of polyamines for productive infection is broadly shared^25^, and targeting polyamine metabolism through host-directed antivirals represents a promising means of blocking virus infection.

## Materials and Methods

### Cell culture

Cells were maintained at 37□C in 5% CO_2_, in Dulbecco’s modified Eagle’s medium (DMEM; Life Technologies) with bovine serum and penicillin-streptomycin. Vero cells (BEI Resources) were supplemented with 10% new-born calf serum (NBCS; Thermo-Fischer) and Huh7 cells, kindly provided by Dr. Susan Uprichard, were supplemented with 10% fetal bovine serum (FBS; Thermo-Fischer).

### Drug treatment

Difluoromethylornithine (DFMO; TargetMol) and N1,N11-Diethylnorspermine (DENSpm; Santa Cruz Biotechnology) were diluted to 100x solution (100mM and 10mM, respectively) in sterile water. For DFMO treatments, cells were trypsinized (Zymo Research) and reseeded with fresh medium supplemented with 2% serum. Following overnight attachment, cells were treated with 100 μM, 500 μM, 1 mM, or 5 mM DFMO. Cells were incubated with DFMO for 96 hours to allow for depletion of polyamines in Huh7 cells. For DENSpm treatment, cells were treated with 100 nM, 1 μM, 10 μM, 100 μM, and 1mM 16 hours prior to infection. During infection, media was cleared and saved from the cells. The same medium containing DFMO and DENSpm was then used to replenish the cells following infection. Cells were incubated at the appropriate temperature for the duration of the infection. Polyamines (Sigma-Aldrich) were added to cells at the time of infection. Methylated spermidine (3-methylspermidine) was derived as described previously^39^ and were added at the time of infection.

### Infection and enumeration of viral titers

RVFV MP-12^40^ and LACV were derived from the first passage of virus in Huh7 cells. CVB3 (Nancy strain) was derived from the first passage of virus in Vero-E6 cells. LACV was obtained from Biodefense and Emerging Infections (BEI) Research Resources. For all infections, DFMO and DENSpm were maintained throughout infection as designated. Viral stocks were maintained at −80□C. For infection, virus was diluted in serum-free DMEM for a multiplicity of infection (MOI) of 0.1 on Huh7 cells, unless otherwise indicated. Viral inoculum was overlain on cells for 10 to 30 minutes, and the cells were washed with PBS before replenishment of media. Dilutions of cell supernatant were prepared in serum-free DMEM and used to inoculate confluent monolayer of Vero cells for 30 min at 37□C. Cells were overlain with 0.8% agarose in DMEM containing 2% NBCS. CVB3 samples were incubated for 2 days, RVFV and LACV samples were incubated for 4 days at 37□C. Following appropriate incubation, cells were fixed with 4% formalin and revealed with crystal violet solution (10% crystal violet; Sigma-Aldrich). Plaques were enumerated and used to back-calculate the number of plaque forming units (pfu) per milliliter of collected volume.

### Virus infectivity assay

RVFV from not-treated or DFMO-treated conditions were incubated at 37^0^C for 24 hours. Subsequent addition of polyamines (10uM spermidine and 10uM spermine) were added to not-treated or DFMO treated virus and incubated at 37^0^C for 24 hours. LACV virus from not-treated or DFMO treated conditions were incubated at 37^0^C for 52 hours. Supernatant was collected at the indicated time points and viral titer was obtained via plaque assay.

### Thin layer chromatography determination of polyamines

Polyamines were separated by thin-layer chromatography as previously described^41^. For all samples, cells were treated as described prior to being trypsinized and centrifuged. Pellets were washed with PBS and then resuspended in 200 uL 2% perchloric acid. Samples were then incubated overnight at 4□C. 200 uL of supernatant was combined with 200 uL 5 mg/ml dansyl chloride (Sigma Aldrich) in acetone and 100 uL saturated sodium bicarbonate. Samples were incubated in the dark overnight at room temperature. Excess dansyl chloride was cleared by incubating the reaction with 100 μL 150 mg/mL proline (Sigma Aldrich). Dansylated polyamines were extracted with 50 μL toluene (Sigma Aldrich) and centrifuged. 5 μL of sample was added in small spots to the TLC plate (silica gel matrix; Sigma Aldrich) and exposed to ascending chromatography with 1:1 cyclohexane: ethylacetate. Plate was dried and visualized via exposure to UV.

### Polyamine luciferase reporter assay

To measure free polyamine levels in cells, a dual-luciferase vector containing the wild-type −1 frameshift antizyme OAZ1 (pC5730) or a dual-luciferase vector containing an in-frame control (pC6154), kindly sent to us by Dr. Tom Dever from the National Institutes of Health, were transfected into cells. Free polyamines modulate OAZ1 mRNA frameshifting and these constructs can measure relative endogenous polyamine concentrations via a dual-luciferase reporter as previously described^42^. 293T cells were seeded with 2% media and drug treated as described above. Cells were transfected with 62.5 ng of either pC5730 or pC6154. After 4h, cells were infected where indicated. After 24 hours of incubation, luminescent signal was measured using the Dual-Luciferase Reporter Assay System (Promega) by measuring both firefly and *Renilla* luciferase with the Veritas Microplate Luminometer (Turner Biosystems). Firefly luciferase was normalized to *Renilla* and the wild-type values were compared to an in-frame control. These values were normalized to untreated cells as relative light units to obtain relative polyamine content.

### RNA purification and cDNA synthesis

Media was cleared from cells and Trizol reagent (Zymo Research) directly added. Lysate was then collected, and RNA was purified according to the manufacturer’s protocol utilizing the Direct-zol RNA Miniprep Plus Kit (Zymo Research). Purified RNA was subsequently used for cDNA synthesis using High Capacity cDNA Reverse Transcription Kits (Thermo-Fischer), according to the manufacturer’s protocol, with 10-100 ng of RNA and random hexamer primers.

### Viral genome quantification

Following cDNA synthesis, qRT-PCR was performed using the QuantStudio3 (Applied Biosystems by Thermo-Fischer) and SYBR green mastermix (DotScientific). Samples were held at 95□C for 2 mins prior to 40 cycles of 95□C for 1s and 60□C for 30s. Primers were verified for linearity using eight-fold serial diluted cDNA and checked for specificity via melt curve analysis following by agarose gel electrophoresis. All samples were used to normalize to total RNA using the ΔC_T_ method. Primers against the RVFV small genome were 5’-CAG-CAG-CAA-CTC-GTG-ATA-GA-3’ (forward) and 5’-CCC-GGA-GGA-TGA-TGA-TGA-AA-3’. Primers for LACV small genome were 5’-GGC-AGG-TGG-AGG-TTA-TCA-AT-3’ (forward) and 5’-AAG-GAC-CCA-TCT-GGC-TAA-ATA-C-3’ (reverse). GAPDH primers were 5’-GAT-TCC-ACC-CAT-GGC-AAA-TTC-3’ (forward) and 5’-CTG-GAA-GAT-GGT-GAT-GGG-ATT-3’ (reverse).

### Genome-to-PFU ratio calculations

The number of viral genomes quantified as described above were divided by the viral titer, as determined by plaque assay, to measure the genome-to-PFU ratio. Values obtained were normalized to untreated conditions to obtain the relative genome-to-PFU ratio.

### Transmission electron microscopy

Four microliters of purified virus were applied to a Formvar-and carbon-coated 400-mesh copper grid (Electron Microscopy Sciences, Hatfield, PA) for 25 seconds. Sample was removed by blotting, and another 4 μL of purified virus was applied for 25 seconds, blotted, and stained with 2% uranyl acetate for 25 seconds. The stained grids were analyzed using a JEOL 1010 transmission electron microscope (Tokyo, Japan) operating at 80 kV. Images were recorded using a Gatan (Pleasanton, CA) UltraScan 4000 charge-coupled-device camera at a magnification of 40,000X.

### Spinoculation and indirect immunofluorescence

Virus was spinoculated onto coverslips by centrifugation at 1200 rpm for 2 hours. Coverslips were subsequently fixed with 4% formalin overnight, washed with PBS, permeabilized and blocked with 0.2% Triton X-100 and 2% BSA in PBS (blocking solution) for 60 minutes at room temperature (RT). Cells were sequentially incubated as follows: Primary mouse anti-Gn antibody (1:1000 in blocking solution, overnight at 4°C), and secondary antibody, goat anti-mouse 488nm, (1:1000 in PBS, 2hr, RT). After washing with PBS, cells were mounted with Everbrite Hardset Mounting Medium (Biotium) overnight. To ensure that signal was not due to impurities, mock-infected supernatant was used as a control and processed in tandem. Samples were imaged with Zeiss Axio Observer 7 with Lumencor Spectra X LED light system and a Hamamatsu Flash 4 camera using appropriate filters using Zen Blue software with a 40X objective (Images were collected with a DeltaVision microscope (Applied Precision) detected with a digital camera (CoolSNAP HQ: Photometrics) with a 60X objective). Images were deconvoluted using SoftWoRx deconvolution software (Applied Precision) and quantified by ImageJ.

### Statistical Analysis

Prism 6 (GraphPad) was used to generate graphs and perform statistical analysis. For all analyses, one-tailed Student’s t test was used to compare groups, unless otherwise noted, with a = 0.05. For tests of sample proportions, p values were derived from calculated Z scores with two tails and α = 0.05.

## Acknowledgments

We are gracious to Thomas Gallagher for critical discussion and helpful insights concerning this project. We thank Susan Uprichard for Huh7 cells and Makio Iwashima for THP-1 cells, as well as helpful discussion. We also thank Shinji Makino and Kaori Terasaki for generously providing the MP-12 strain of RVFV. We appreciate immunofluorescent imaging help and input from the labs of Ivana Kuo and Jordan Beach. Synthesis of (*R*)-isomer of 3-MeSpd was supported by grant of Russian Scientific Foundation #17-74-20049.

